# Evaluation of GABA_A_R-mediated inhibition in the human brain using TMS-evoked potentials

**DOI:** 10.1101/2022.10.31.514225

**Authors:** Dominika Šulcová, Adriana Salatino, Adrian Ivanoiu, André Mouraux

## Abstract

GABA_A_ receptor (GABA_A_R) – mediated inhibition participates in the control of cortical excitability, and its impairment likely contributes to the pathologic excitability changes that have been associated with multiple neurological disorders. Therefore, there is a need for its direct evaluation in the human brain, and the combination of transcranial magnetic stimulation (TMS) and electroencephalography (EEG) might represent the optimal tool. TMS-evoked brain potentials (TEPs) capture the spread of activity across the stimulated brain network, and since this process at least partially depends on the GABA_A_R-mediated inhibition, TEPs may constitute relevant biomarkers of local GABA_A_ergic function.

Here, we aimed to assess the effect of GABA_A_Rs activation using TEPs, and to identify TEP components that are sensitive to the state of GABA_A_ergic inhibition. In 20 healthy subjects, we recorded TEPs evoked by sub- and supra-threshold stimulation of the primary motor cortex (M1), motor-evoked potentials (MEPs) and resting-state EEG (RS-EEG). GABA_A_Rs were activated (1) pharmacologically by oral administration of alprazolam compared to placebo within each subject, and (2) physiologically using a sub-threshold conditioning stimulus to characterize the effect of short-latency intracortical inhibition (SICI).

In supra-threshold TEPs, alprazolam suppressed the amplitude of components N17, N100 and P180, and increased component N45. The pharmacological modulation of N17 correlated with the change observed in MEPs and with the alprazolam-induced increase of lower β-band RS-EEG. Only a reduction of N100 and P180 was found in sub-threshold TEPs. TEP SICI manifested as a reduction of N17, P60 and N100, and its effect on N17 correlated with the alprazolam-induced N17 suppression and β increase. Our results indicate that N17 of supra-threshold TEPs could serve as a non-invasive biomarker of local cortical excitability reflecting the state of GABA_A_R-mediated inhibition in the sensorimotor network. Furthermore, the alprazolam-induced increase of β-band oscillations possibly corresponds to the increased inhibitory neurotransmission within this network.

## 1. Introduction

Fine tuning of the activity of excitatory and inhibitory systems is crucial for maintaining physiological brain functions [1, 2], its dysregulation potentially leading to pathological changes in cortical excitability. In various neurological disorders, including epilepsy [3], schizophrenia [4], amyotrophic lateral sclerosis (ALS) [5] and Alzheimer’s disease (AD) [6], the balance is tipped towards increased excitation that can be associated with excitotoxicity. While the original cause of such disbalance is difficult to pinpoint, the suspicion often falls on impaired inhibition. Inhibitory neurons represent only a small portion of all cortical neurons (10-15% reported in rodents; [7]). However, they come in a great variety of morphologic and connectivity types [8, 9], forming a complex and flexible system holding control over principal excitatory neurons and regulating the dynamics of entire brain networks [10-12]. For this reason, minor functional impairment of inhibitory neurons could produce extensive changes in brain function.

The most common inhibitory neurotransmitter of the brain is gamma-butyric acid (GABA). Release of GABA by GABAergic inhibitory interneurons exerts fast inhibitory effects via postsynaptic activation of GABA_A_ receptors (GABA_A_Rs) that react to the neurotransmitter binding by increasing the membrane permeability for chloride ions [13]. Resulting hyperpolarization of the postsynaptic membrane gives rise to an inhibitory potential at the target neuron that contributes to the synchronisation of the network oscillatory activity [10]. The strength of GABA_A_R-mediated inhibition and therefore the performance of the inhibitory system can be influenced at multiple levels both pre- and post-synaptically, with any change triggering a cascade of compensatory or adaptive processes. Accordingly, neurological disorders that are characterized by an excitation-inhibition imbalance towards reduced inhibition have been previously linked both to the malfunction of inhibitory interneurons [14-16] and to alternations in GABA_A_R expression and function [17-20].

Considering the possible involvement of GABAergic inhibitory dysfunction in a wide array of neurological diseases, it is important to establish reliable methods to assess the functional state of GABA_A_R-mediated neurotransmission in humans. This might be particularly useful for neurodegenerative disorders such as AD or ALS, where impairment of inhibitory neurotransmission is likely to occur prior to clinical symptoms and where early diagnosis and early treatment could positively influence the prognosis. For this purpose, the combination of transcranial magnetic stimulation (TMS) and electroencephalography (EEG) represents a promising tool because it allows to directly probe the functional properties of cortical networks. TMS can non-invasively and painlessly stimulate a precisely defined cortical area [21, 22], and concomitant EEG can capture in real time the spread of the TMS-evoked activity across the brain [23, 24]. Importantly, both methods are long established in experimental and clinical research, with few contra-indications and relatively low cost, which favours them for routine application in large patient groups.

TMS combined with EEG has already been successfully used to evaluate the state of inhibition in the human brain by characterizing the changes in TMS-evoked brain potentials (TEPs) induced by the experimental activation of GABA_A_Rs. This can be achieved by different means. Receptor function can be modulated pharmacologically with benzodiazepines that enhance GABA_A_R-mediated inhibition globally, as they act on selected subtypes of the receptor across the whole nervous system. Using this approach, Premoli et al. in their pioneer study identified specific TEP components sensitive to GABA_A_R activation [25]. Alternatively, it is possible to trigger GABA_A_R-mediated inhibition locally by pre-activating inhibitory interneurons at the site of stimulation with a sub-threshold conditioning TMS pulse. This was originally described in motor-evoked potentials (MEPs) using the so-called “paired-pulse paradigm” in which two TMS pulses are delivered in rapid succession over the primary motor cortex (M1), a sub-threshold conditioning TMS pulse followed by a supra-threshold TMS pulse. As compared to the amplitude of the MEP elicited by a single supra-threshold TMS pulse, the amplitude of the MEP elicited by paired pulses is markedly reduced, and this effect has been attributed to the triggering of GABA_A_R-mediated short-latency intracortical inhibition (SICI) [26, 27]. More recently, several groups described SICI also in M1 TEPs, yet their findings were highly variable as well as the methodology they used [28-33].

In the present study, we aimed to identify TEP-based biomarkers of GABA_A_R-mediated inhibition in the human brain and to verify its reliability using different approaches. First, we activated the inhibitory system globally using alprazolam, a classic benzodiazepine that acts on GABA_A_Rs containing α1, α2, α3 and α5 subunits [34]. We described its effect on TEPs evoked by stimulation of the left M1 using both sub- and supra-threshold stimulation intensities by comparing the effects of alprazolam to those of an active placebo. In addition, we recorded resting state EEG to extract previously established biomarkers of benzodiazepine-induced sedation and related them to changes observed in TEPs. Second, we used paired-pulse stimulation to activate local inhibitory interneurons and characterized the effect of SICI in M1 TEPs. We assumed that the increase of GABA_A_R-mediated neurotransmission produced by the two methods would result in similar changes for M1 TEP components that are specifically dependent on the state of GABA_A_R neurotransmission locally within the stimulated cortex, whereas alprazolam could produce additional changes in M1 TEP components due to its inhibitory effect in the entire brain.

## 2. Materials and Methods

### 2.1. Ethical approval

All experiments were conducted according to the latest revision of the Declaration of Helsinki. The protocol and all procedures were approved by the local Ethical Committee. All participants were thoroughly informed of the protocol and applied procedures, gave a written informed consent before taking part in the study, and were financially compensated for their participation.

### 2.2. Data and code availability

The data presented in this study are not freely available, because the informed consent signed by the participants did not include the agreement to public data sharing. However, it is possible to request the raw or pre-processed datasets from the corresponding author, under the condition of an established inter-institutional agreement of the data transfer. MATLAB and R scripts used to process and evaluate the data are accessible from the GitHub page of the corresponding author (https://github.com/DominikaSulcova/GABA-AD), and the letswave6 toolbox that provided called functions can be downloaded here: https://letswave.org.

### 2.3. Participants

20 healthy volunteers (10 males, mean age 24) took part in the study. All participants filled a detailed questionnaire and undertook a brief neurological examination to exclude candidates with any contraindication for experimental procedures, such as epilepsy (or familiar history thereof), migraine, presence of a metal object in the body (e.g. insulin pump, pacemaker, metal prothesis), confirmed neurological pathology, or use of medication known to alter cortical excitability [35].

### 2.4. Experimental design

Two experimental sessions, each lasting approximately 5.5 h and separated by at least one week, were conducted at the premises of the UCLouvain Institute of Neuroscience (IoNS). The experiments always started in the morning at times as similar as possible in both sessions for each subject. The subjects were asked to sleep sufficiently, to refrain from consumption of alcohol the preceding evening and to avoid caffeinated beverages in the morning. Each session consisted of the baseline recording of resting state EEG (RS-EEG) and TMS stimulation of the left (dominant) primary motor cortex (M1) combined with EEG for the recording of TEPs and EMG for the recording of MEPs. An additional block of TMS stimulation of the left angular gyrus was performed but the results are not discussed in this publication (see Supplementary materials). Subjects then received a single dose of either alprazolam (Xanax, 1mg) or active placebo (Zyrtec, 10 mg), and the recording was repeated starting 1.5 h after the medication. This timing corresponds to the average peak levels of alprazolam in the blood following oral administration [36, 37]. The order of the two sessions was balanced across subjects. The medication was provided to the participant by a third party so that both subjects and experimenters performing the experimental sessions and analysing the recorded data were blinded to the order of the medication.

### 2.5. EEG recording

The EEG was recorded using the NeurOne EEG system (Bittium NeurOne Tesla; Bittium Corporation, Oulu, Finland) and a 32 channel EEG cap mounted with TMS-compatible Ag/AgCl electrodes (EasyCap GmbH, Herrsching, Germany). The signal was recorded at a 20 kHz sampling rate with a 5000 Hz low-pass filter and a DC filter. The averaged signal from both mastoids was used as reference, an extra ground electrode was placed on the forehead. Electrode impedances were monitored during the session and kept below 5kΩ. Two blocks of RS-EEG with eyes open and closed, each lasting at least 3 min, were recorded at baseline and after the medication, before TMS stimulation. During the recording with eyes open, subjects kept their gaze on a stable point placed in such way that its fixation did not require any muscular effort.

A thin plastic film was placed over the electrodes and subjects wore an extra rubber cap on top of the plastic film to minimize artifacts caused by direct contact of electrodes with the TMS coil. These additional layers also helped to reduce bone conduction of the noise generated by TMS stimulation. In addition, subjects listened to a masking noise created from an original recording of the TMS click to further reduce possible auditory artifacts [38].

### 2.6. EMG recording

Disposable surface gel electrodes were placed over the first dorsal interosseous muscle (FDI) of the right hand using the belly-tendon montage, with the ground electrode placed at the right wrist over the styloid process of radius. Motor-evoked potentials (MEPs) were recorded at a 1,024 Hz sampling rate using the MOBI amplifier (TMSi MOBI; Twente Medical Systems International B.V., Oldenzaal, The Nederlands).

### 2.7. TMS stimulation and neuronavigation

A 3D T1-weighted structural magnetic resonance image (MRI) of each participant’s whole brain was acquired beforehand at the Department of radiology of the Cliniques universitaires Saint-Luc (1×1×1 mm; 3 T Achieva; Philips Healthcare, Amsterdam, The Netherlands). The Visor2 neuronavigation system (Visor 2.3.3; Advanced Neuro Technologies, Enschede, The Netherlands) was used to verify the accurate placement of the TMS coil and to monitor its position throughout the experiment. A 3D model of the scalp and brain surface was reconstructed using the individual MRI data and co-registered with the real space using landmark-based markers (nasion and tragi) and head-shape matching [39].The position of the head as well as the TMS coil was continually registered with the Polaris infrared optical tracking system (Polaris Spectra; Northern Digital Inc. Europe, Radolfzell, Germany).

Biphasic TMS pulses were delivered manually using a MagPro X100 magnetic stimulator (MagPro X100 with MagOption; MagVenture, Farum, Denmark). A figure-of-eight TMS coil with an outer diameter of 75 mm (C-B60; MagVenture, Farum, Denmark) was tangentially placed on the scalp and centred over the target in the left M1 (Fig 1A). Target location was determined functionally as the site eliciting the largest TMS-evoked MEP in the relaxed right FDI when stimulating with slightly sub-threshold intensity and inducing electrical currents in the brain with an anterior–posterior posterior– anterior direction. The optimal coil orientation was adjusted individually, with the handle always pointing laterally and posteriorly at an approximate 45° angle from the midline. Target parameters were saved in the neuronavigation system to keep stimulation sites identical in the second experimental session.

**Fig 1.**
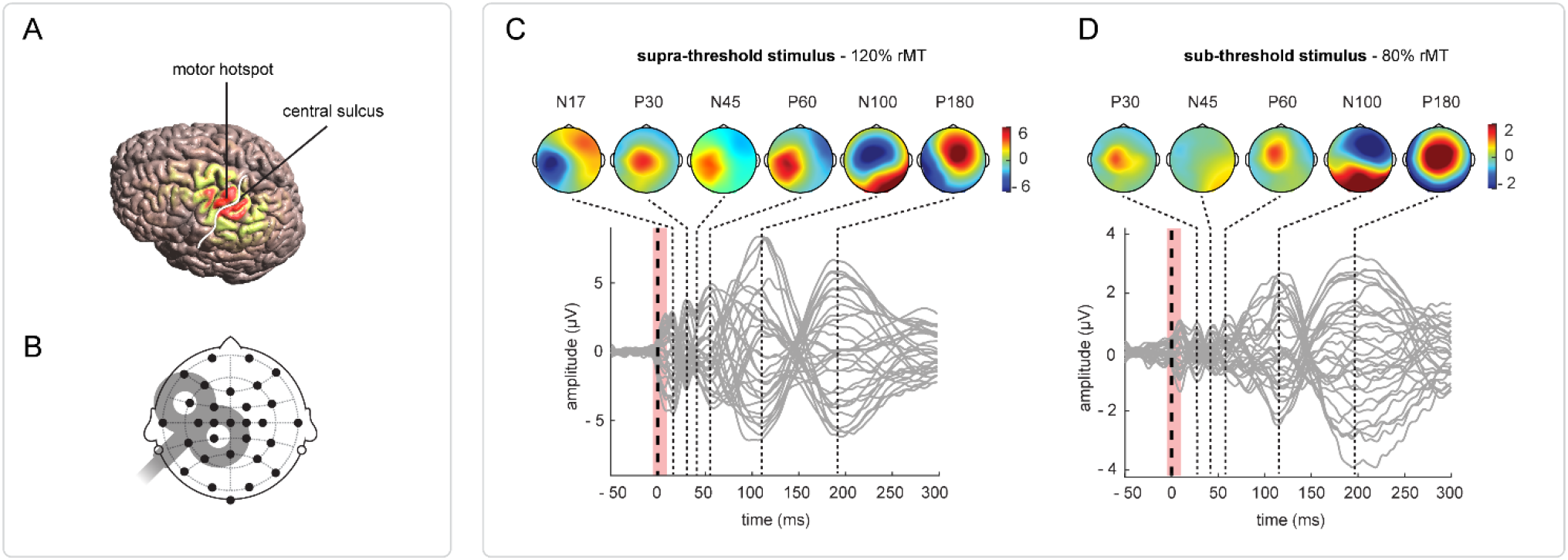
TMS-EEG over the primary motor cortex. Targeting M1: (A) Magnetic field spreading across the brain surface when targeting the hand area of the left primary motor cortex with TMS (modelled on the default brain with the simulation software *simNIBS* [97]).(B) Approximate placement of the TMS coil over the head. Black circles represent EEG electrodes in the used setup. (C-D) **Baseline M1 TEPs** evoked by supra-threshold (C) and sub-threshold (D) M1 stimulation. Grey traces represent the signal of all electrodes recorded at the baseline in the placebo sessions (there were no substantial differences between baseline TEPs of both sessions), TMS stimulus is marked by a thick dashed line, the interval interpolated following the removal of TMS artifact is shaded in pink. Dotted lines mark peak latencies of TEP components, associated topographies are shown above.

The resting motor threshold (rMT) was set to the minimal stimulation intensity eliciting a motor response with the amplitude of least 50 µV in at least 5 trials out of 10 [40]. Its value was verified and adjusted, if necessary, before the data acquisition after medication, in order to record reliable MEPs. The mean rMT at baseline (in % of the maximum stimulator output ± SEM) was 49.3 ±1.4 in the placebo session and 49.8 ±1.4 in the alprazolam session. This difference was not significant (repeated measures ANOVA: F(1, 19) = 1.05, p = 0.32).

Three types of TMS stimuli were applied over M1: supra-threshold single pulses at 120% rMT (testing stimulus, TS); sub-threshold single pulses at 80% rMT (conditioning stimulus, CS); and paired pulse TMS consisting of a CS followed by a TS with an inter-pulse interval of 2.5 ms (CS-TS). 60 stimuli per stimulation type were delivered to M1 in a semi-random order (maximum 3 repetitions of the same condition) and a randomly varying inter-trial interval (4-6 s). The sequence was split into three blocks of 60 stimuli with simultaneous EEG and EMG recordings, separated by a short break.

### 2.8. Data processing

TEPs, MEPs and RS-EEG were analysed offline in Matlab (MathWorks, Inc., Natick, Massachusetts, United States) using Letswave 6 (an open-source EEG signal processing toolbox, https://www.letswave.org/) and custom scripts.

#### TEPs

Data pre-processing generally followed a previously established pipeline [41]. The EEG signals were first re-referenced to the common average of all scalp channels. The DC shift was removed and a linear detrend was applied. The continuous EEG recordings were then epoched around the onset of the TMS pulses from –1000 ms to +2000 ms. The large-amplitude artifact caused by the TMS pulse was removed using cubic interpolation (from -5 to +10 ms) and the signals were down-sampled to 2 kHz. A first round of ICA was performed to remove the remaining tail of the TMS-evoked muscle artifact, because the sharp edge of the tail might otherwise interfere with applied frequency filters. Following this ICA, the data was bandpass filtered between 0.1 and 80 Hz (Butterworth, 4^th^ order) and notch filtered at 50 Hz (FFT linear filter, notch width 2 Hz, slope cut-off width 2 Hz). In the next step, data were visually inspected to remove epochs containing excessive noise or isolated outbursts thereof. Any trials associated with a missed TMS stimulation (in case of an unexpected head movement causing coil misplacement, etc) were discarded. The average number of remaining epochs (± SD) was 48.0 ± 6.9 in the placebo session and 47.8 ± 6.9 in the alprazolam session. The second round of ICA was used to remove any remaining artifacts such as eye blinks, horizontal eye movements, persistent muscle activity and electrode noise. Artefactual components were first identified automatically with the Multiple Artifact Rejection Algorithm (MARA), a linear classifier originally trained on adult EEG data [42]. Pre-selected components were then manually assessed based on their topography, time course and power spectrum density to remove only components with clear non-neural origin (a detailed characterization of different types of removed artifacts is available in Supplementary materials). All epochs were baseline corrected to the interval from -200 to -5 ms and the trials were averaged for each subject/condition.

A time window between 10 and 300 ms post stimulus was considered for the analysis. Peak latencies of TEP components were identified in the temporal profile of the Global Field Power (GFP) calculated from the baseline recordings according to the following equation:

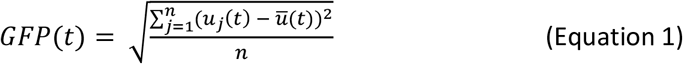

where *t* corresponds to a timepoint, *n* is the number of electrodes, *u*_*j*_ represents the voltage measured at each electrode and *ū* the mean voltage across all electrodes. Identified TEP components were then named according to their polarity as seen in the signal at electrode Cz (positive = P, negative = N) and their average latency in ms. Based on the results of the topographical analysis performed on baseline recordings [43], we identified three electrodes of interest (EOIs) for each TEP component and each type of stimulus (expectedly, the EOIs were the same for supra-threshold TMS pulses delivered over M1 in the conditions TS and CS-TS). The peak amplitude of each TEP component was extracted at the subject level from the averaged signals measured at the EOIs using a semi-automatic approach (the process is described in more detail in the Supplementary materials). This approach allowed to account for individual differences in the latency of TEP components while keeping other extraction parameters constant across subjects.

#### MEPs

Raw EMG signals were corrected for DC shift, linear detrended and high-pass filtered with 4 Hz cut-off (Butterworth, 4^th^ order). The signals were then segmented around the TMS pulse into epochs ranging from -100 to +400 ms. Epochs with excessive baseline activity were identified and discarded using an automated selection process based on the baseline mean root square (MRS) value (see Supplementary materials), with 56.5 ± 3.4 (SD) epochs retained on average across datasets. The signals were then baseline corrected by subtracting the mean amplitude of the interval between -200 and 0 ms. The peak-to-peak MEP amplitude was extracted for further analysis.

#### RS-EEG

Raw RS-EEG recordings were down-sampled to 2 kHz, re-referenced to a common average of all scalp channels and bandpass filtered between 0.1 and 45 Hz (Butterworth, 4^th^ order). Following a visual inspection, 2 minutes of clean signal were cut out from each of the two original 3-minute recordings (eyes open and closed). An Independent Component Analysis (ICA; Infomax algorithm), applied on the concatenated EEG signals, was then used to remove artifacts with clear non-neural origin (e.g. tonic muscle contraction, eyeblinks, electrode noise). Then, both ICA-denoised datasets were segmented in 50% overlapping epochs lasting 12 s for the analysis of δ band oscillations and 4 s for the analysis of higher frequency bands. A Hanning window was applied to each epoch to minimize spectral leakage. The epochs were transformed using the Fast Fourier Transform and averaged across epochs to express mean amplitude for each frequency bin between 0.1 and 45 Hz. EEG channels were clustered into five regions of interest (ROIs; frontal: Fp1, Fp2, Fz, F3, F4, F7, F8; central: FC1, FC2, Cz, C1, C2, CP1, CP2; left temporal: FC5, T7, C3, CP5; right temporal: FC6, T8, C4, CP6; occipital: P3, P4, Pz, P7, P8, O1, O2).

Baseline data of each subject were used to identify the Individual Alpha peak Frequency (IAF) and the Transition Frequency (TF, frequency marking the transition between theta band oscillations and the alpha peak) as described in [44]. Based on these values, the frequency spectra were split into the following bands: δ (0.1Hz – [TF – 4Hz]), θ ([TF – 4Hz] – TF), lower α1 (TF - [TF + (IAF -TF)/2]), lower α2 ([TF + (IAF -TF)/2] - IAF), higher α3 (IAF - [IAF + (IAF -TF)/2]), lower β1 ([IAF + (IAF -TF)/2] – 20Hz), higher β2 (20 – 30Hz), and γ (30 – 45Hz).

Three RS-EEG measures were selected for further statistical analyses: (1) The average amplitude of lower β frequency band was extracted from the central ROI in the ‘eyes-open’ dataset and used to characterize the effect of alprazolam, because an increase in lower β oscillations following the administration of benzodiazepines was previously reported in multiple studies and proposed as a RS-EEG biomarker of the benzodiazepine effect [45-48]. (2) The Alpha Attenuation Coefficient (AAC) was calculated as the ratio of the broadband α amplitude in ‘eyes closed’/’eyes open’ over the occipital ROI. The AAC was previously established as a measure of drowsiness based on the observation that with increasing subjective drowsiness the α amplitude increases with eyes open and decreases with eyes closed [49, 50]. Similar changes in α oscillations were also observed after medication with benzodiazepines [51-53]. Therefore, we expected a decrease in AAC following the administration of alprazolam. (3) The Spectral Exponent (SE) was calculated as the slope of the 1/f-like non-oscillatory spectral component under the log-log transform across the lower part of the spectrum (δ – lower β) using the data averaged across all ROIs (for a detailed description see [54]). SE was previously linked to the excitation-inhibition balance in neural circuits [55] and its increased negativity was observed during states of reduced consciousness [54]. Based on these reports, we expected the SE to gain more negative values following the administration of alprazolam.

### 2.9. Statistical analysis

Statistical analyses were performed using Matlab and R [56]. A Linear Mixed Model (LMM) analysis was implemented and interpreted using R packages *lme4* [57] and *lmerTest* [58].

The effect of alprazolam was evaluated in relation to single-pulse TEPs, MEPs, RS-EEG, as well as the extent of SICI in TEPs and MEPs. The post-medication change was calculated as percentage of the post-medication value relative to the baseline value for RS-EEG and MEPs, and as the difference between baseline and post-medication measures for TEP amplitudes. Each obtained outcome variable was then separately evaluated using a LMM model that allowed for a random intercept (Equation 2). The predictor of interest was the factor Medication with 2 levels (placebo, alprazolam). When evaluating alprazolam-induced change in TEPs, the change in rMT was entered as a time-varying covariate after being centred to mean 0. Final p value was obtained using the Kenward-Roger approximation.

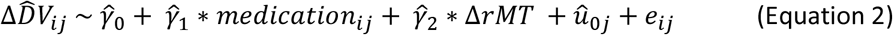

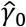represents the common parameter for the intercept, 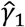the common parameter for the main predictor, *û*_0*j*_ stands for the individual variability in the intercept (= random effect), and *e*_*ij*_ for the residual error.

For each subject, short latency intracortical inhibition (SICI) over M1 was quantified in both sessions at baseline, i.e. before medication. To assess the effect of the paired pulse stimulation on the amplitude of MEPs and individual TEP components, a LMM model with random intercept was used (Equation 3). The measured raw amplitude was entered as the dependent variable, the main predictor was the factor Stimulus with 2 levels (TS, CS-TS) and the session ID was entered as a covariate. Final p value was obtained using the Kenward-Roger approximation.

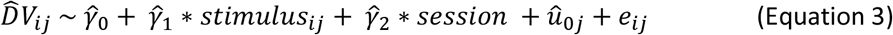

For the purposes of visualization and correlational analysis, SICI of MEPs was expressed as the percent change of peak-to-peak MEP amplitude elicited by CS-TS compared to the MEP evoked by the TS stimulus alone, while SICI of each TEP component was computed as a simple difference between the amplitude of CS-TS TEPs and TS TEPs. To evaluate possible correlations between the selected variables, Pearson correlation coefficients were computed and tested against 0, with the significance level adjusted using Bonferroni correction in case of multiple comparisons. In certain cases, if the preliminary visualization suggested a non-linear relationship between variables, the data were ranked, and a Spearman correlation coefficient was used instead.

## 3. Results

All mean values are reported ± standard error of mean (SEM).

### 3.1. TEP features depend on TMS stimulation intensity

Supra-threshold TMS stimulation of M1 evoked consistent TEPs composed of 6 components that showed latencies similar to those previously reported in the literature: N17, P30, N45, P60, N100 and P180 (Fig 1C). The components were clearly identified as local maxima in the GFP of the group average TEP waveform, except for N45 that might have been overlayed by the peak corresponding to P60. By contrast, only five TEP components were identified in TEPs evoked by sub-threshold TMS stimulation of M1 at latencies corresponding to P30, N45, P60, N100 and P180 (the earlier N17 component was impossible to distinguish in most individual datasets; Fig 1D). TEPs elicited by sub-threshold stimulation had a lower amplitude and the topography of all components (with the exception of P30) differed substantially from those evoked by supra-threshold stimulation. Baseline M1 TEPs were analysed in detail and compared with TEPs elicited by stimulation of the angular gyrus in a separate publication [43].

### 3.2. Alprazolam modulates peak amplitude of specific TEP components

The peak amplitude of individual TEP components served as the main outcome variable when evaluating the effect of medication on TEPs. Mean observed changes and the outcome of the statistical analysis are summarized in Table 1. When considering the TEPs evoked by supra-threshold M1 stimulation (Fig 2A,B), an opposite effect of medication was observed for N17 amplitude: while alprazolam led to a decrease of this component, placebo caused an increase. Subsequent TEP components were only minimally influenced by placebo, whereas alprazolam caused an increase of N45 and a reduction of N100 and P180. In addition, a borderline significant reduction caused by alprazolam was found in P60 (p = 0.05).

**Fig 2.**
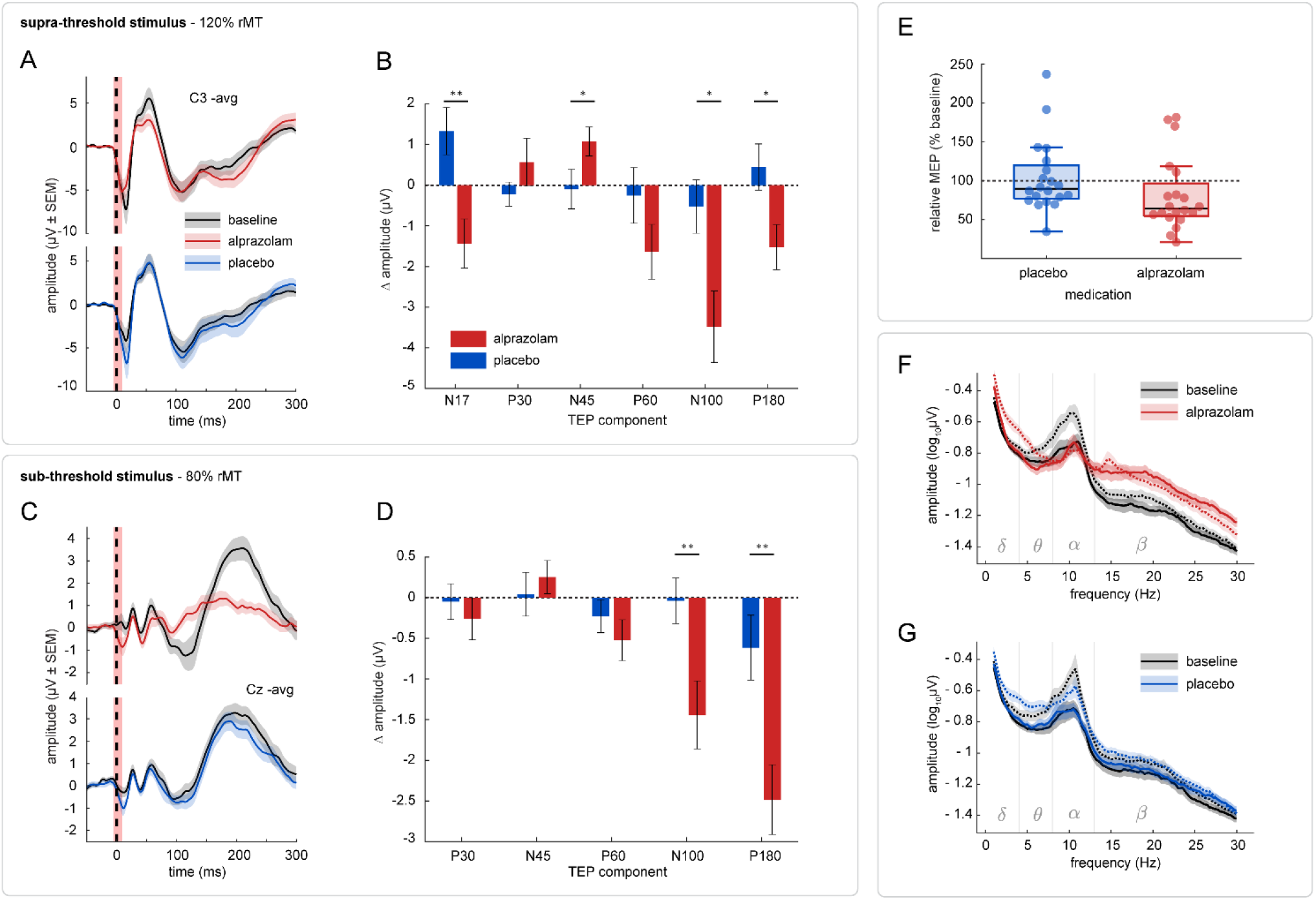
Effect of alprazolam. TEPs: (A, C) Group average signals from the most representative electrode for supra- and sub-threshold M1 TEPs are plotted at the baseline and after the medication (baseline: black, post alprazolam: red, post placebo: blue). Shaded areas represent the SEM. (B, D) Average modulations of peak amplitudes (± SEM) in supra- and sub-threshold M1 TEPs are illustrated with a bar plot (alprazolam: red, placebo: blue). Positive and negative values correspond to an increase and decrease of amplitude, respectively. In case of negative components, the original voltage measures were flipped to better convey the change induced by the medication. Asterisks denote the significance of the difference between the two sessions (* = p < 0.05; ** = p < 0.01). **MEPs:** (E) Boxplots show the modulation of peak-to-peak MEP amplitude evoked by supra-threshold stimulation of M1, values are expressed in % of baseline (alprazolam: red, placebo: blue). No significant effect of medication was found. **RS-EEG:** (F, G) Frequency spectra of RS-EEG recordings (central region) from the alprazolam and placebo session are visualized as the log_10_ of the average amplitude at frequencies 1 – 30Hz (baseline: black, post-alprazolam: red, post-placebo: blue). Full lines represent the signal with eyes open, dotted lines the signal with eyes closed. Shaded areas mark the SEM.

**Table 1.** Mean change in TEP amplitude and corresponding statistics. Amplitude change was computed as the difference between measurement post-medication and the baseline, in the table represented in μV ± SEM. Positive numbers mark an increase of the amplitude and vice versa, in case of negative components original voltage measures were multiplied by -1.

| stimulus | peak | mean change in amplitude |  | F statistic | df | p value |  |
| --- | --- | --- | --- | --- | --- | --- | --- |
|  |  | placebo | alprazolam |  |  |  |  |
| M1<br>120 %rMT | N17 | 1.33 ± 0.58 | - 1.44 ± 0.60 | 10.62 | 1, 18.54 | < 0.01 | ** |
|  | P30 | - 0.22 ± 0.30 | 0.57 ± 0.59 | 1.77 | 1, 18.50 | *n.s.* |  |
|  | N45 | - 0.10 ± 0.49 | 1.08 ± 0.35 | 4.91 | 1, 18.39 | < 0.05 | * |
|  | P60 | - 0.25 ± 0.69 | - 1.64 ± 0.68 | 4.43 | 1, 17.55 | 0.05 |  |
|  | N100 | - 0.52 ± 0.66 | - 3.48 ± 0.88 | 7.95 | 1, 18.50 | < 0.05 | * |
|  | P180 | 0.45 ± 0.57 | - 1.52 ± 0.55 | 6.19 | 1, 18.53 | < 0.05 | * |
| M1<br>80 %rMT | P30 | - 0.05 ± 0.22 | - 0.26 ± 0.26 | 0.40 | 1, 18.52 | *n.s.* |  |
|  | N45 | 0.04 ± 0.27 | 0.25 ± 0.20 | 0.00 | 1, 17.54 | *n.s.* |  |
|  | P60 | - 0.23 ± 0.20 | - 0.52 ± 0.25 | 0.96 | 1, 18.46 | *n.s.* |  |
|  | N100 | - 0.04 ± 0.28 | - 1.44 ± 0.42 | 11.19 | 1, 18.35 | < 0.01 | ** |
|  | P180 | - 0.61 ± 0.40 | - 2.48 ± 0.43 | 10.00 | 1, 18.54 | < 0.01 | ** |

In TEPs elicited by sub-threshold M1 (Fig 2C,D), late TEP components N100 and P180 were clearly suppressed by alprazolam as compared to placebo, while earlier components showed no significant change in amplitude.

### 3.3. Drug-induced change in MEP amplitude is correlated with change in amplitude of TEP N17

When evaluating the effect of medication on the amplitude of single-pulse MEPs, we found that while alprazolam caused an average decrease by 18.8 ± 10.6 %, placebo led to an average increase by 3.8 ± 10.3 % (Fig 2E). However, the alprazolam-induced change was not significantly different from placebo (F_1, 18.54_ = 3.8, p = 0.07).

We further explored the relationship between the drug-induced changes in MEP and TEP amplitudes. Since the preliminary data visualization suggested possible linear relationships between MEP and TEP changes, we calculated Pearson’s correlation coefficients for MEPs and the 6 components of supra-threshold M1 TEPs (Bonferroni corrected). From all tested components, only N17 showed a significant correlation between both variables when data from both sessions were pooled together (r(38) = 0.46, t = 3.16, p < 0.01, adjusted R^2^ = 0.19; see Fig 4A). This correlation remained significant and showed very similar slopes when the sessions were considered separately. This indicates that the modulation of N17 amplitude is directly associated with the modulation of cortical motor output.

### 3.4. The effect of alprazolam manifests in resting state EEG

Alprazolam caused multiple changes in the oscillatory content of the RS-EEG. We observed an overall increase in δ band amplitude, an increase in the amplitude within the high α and both β bands in the eyes open condition, and a substantial suppression of oscillations across all α sub-bands in the eyes closed conditions (Fig 2F,G). For all three RS-EEG measures that were selected for the statistical evaluation, we observed a change in the same direction following the administration of both drugs, which was significantly greater after alprazolam compared to placebo. Mean observed changes as well as the outcome of the statistical analysis are summarized in Table 2. As expected, the β increase over the central region proved to be a particularly consistent marker of the effect of alprazolam. The AAC decrease was mostly related to an extensive suppression of the α peak in the eyes closed condition. Lastly, the SE became more negative after alprazolam, which corresponds to a steeper slope of the decay in the power spectrum density (PSD).

**Table 2.**
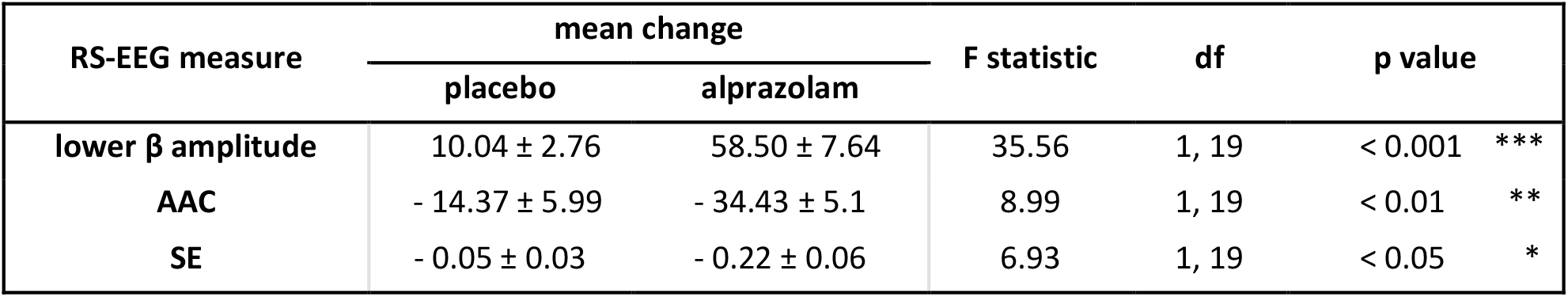
Mean change in RS-EEG measures and corresponding statistics. The change in lower β amplitude and in the AAC is represented in % of baseline ± SEM. The SE change was computed as the difference between the measurement post-medication and the baseline.

### 3.5. Alprazolam-induced reduction of N17 correlates with the increase in β band amplitude

To further investigate the connection between the TEP component N17 and the state of GABA_A_ergic inhibitory system, we explored the relationship between the medication-induced change in N17 and the established RS-EEG markers. From the three tested RS-EEG measures, only the change in low β amplitude proved to be significantly correlated with the change in N17 (Fig 4B). Moreover, we observed a different relationship between the two variables depending on the medication – while placebo showed no significant association (r(18) = 0.01, t = 0.06, adjusted R^2^ = -0.06), a clear negative correlation was found in measures from the alprazolam session (r(18) = -0.6, t = -3.21, p < 0.01, adjusted R^2^ = 0.39): greater increases in β were associated with a stronger decrease of the N17.

### 3.6. SICI leads to a selective suppression of M1 TEP components

Short-latency intracortical inhibition (SICI) in the M1 was recorded before medication in both sessions using a paired-pulse TMS protocol. For this purpose, a sub-threshold conditioning stimulus (CS, 80 %rMT) was applied 2.5 ms before the supra-threshold testing stimulus (TS, 120 %rMT), and resulting MEPs and TEPs (CS-TS) were compared to measures evoked by the TS alone. As expected, SICI manifested as a reduction of the MEP peak-to-peak amplitude (Fig 3C) that was suppressed in average by 74.94 ± 5.2 % in the baseline recording of the placebo session and by 75.21 ± 5.51 % in the baseline recording of the alprazolam session. In the M1 TEPs, a significant reduction of amplitude was observed for components N17 (F_1,58_ = 39.37, p < 001), P60 (F_1,58_ = 12.98, p < 001) and N100 (F_1,58_ = 10.7, p < 01), and these changes were consistent across both baseline recordings (Fig 3A,B). Mean amplitude modulation of all TEP components and the outcome of the statistical analysis are summarized in the Table 3.

**Fig 3.**
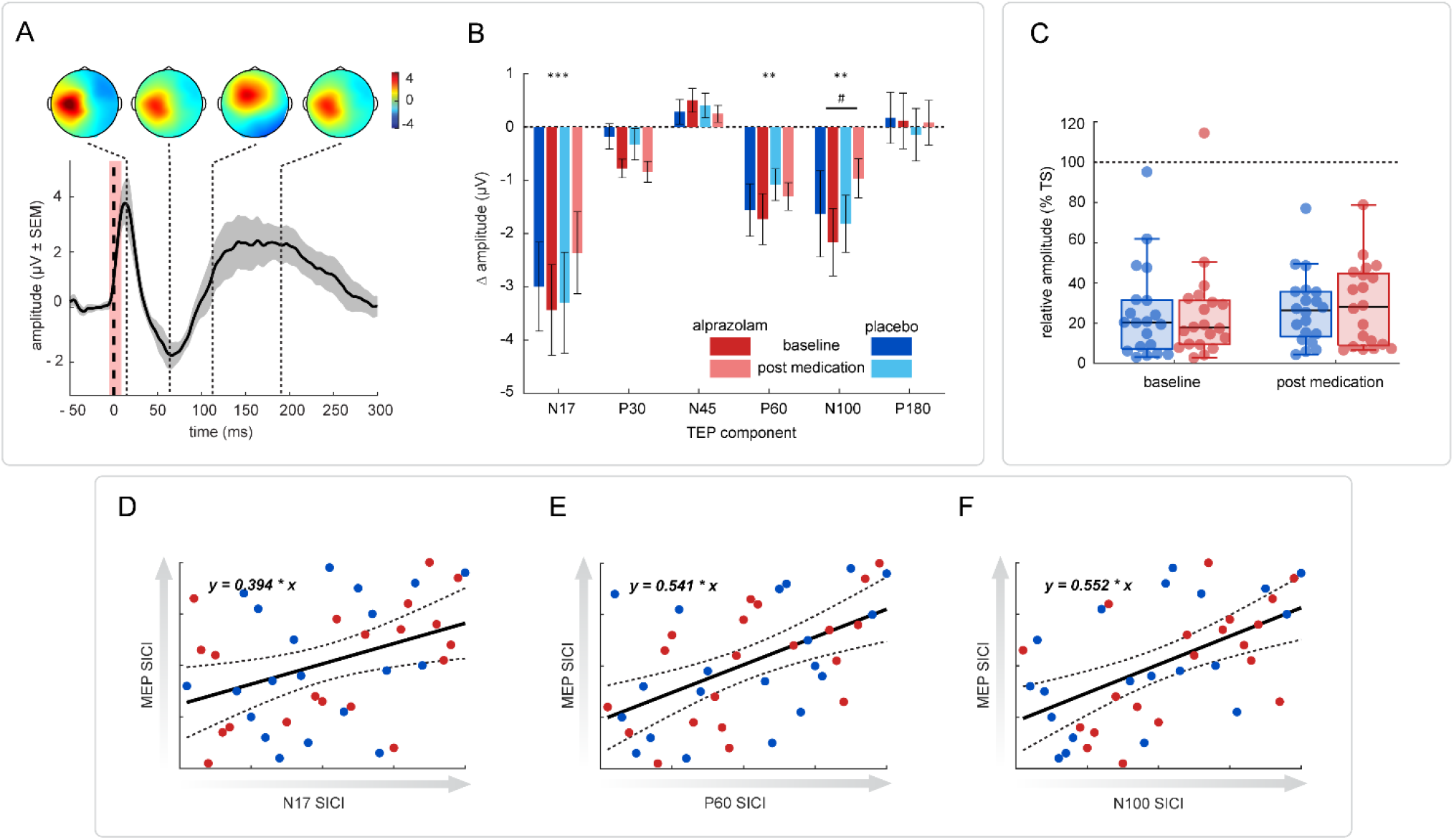
Effect of SICI. TEPs: (A) The time-course and topographic distribution of TEP SICI was obtained by point-by-point subtraction of the single-pulse TEP from the paired-pulse TEP over the whole explored time-window. The signal from C3 electrode is shown, the grey shaded area represents the SEM. The thick dashed line marks the TMS stimulus, the shaded pink area corresponds to the interval interpolated following the TMS artifact removal. Dotted lines mark latencies at which distinct topographical distributions of voltage change could be identified, as illustrated in the upper row. (B) The extent of TEP SICI at each TEP component was calculated as a subtraction of the TS TEP amplitude from the CS-TS TEP amplitude. The bar graph shows the average effect of SICI (μV ± SEM) separately for both experimental sessions at the baseline (dark colours) and following the medication (light colours; alprazolam: red, placebo: blue). In case of negative components, the original voltage measures were flipped to better convey the direction of change. Asterisks denote the significance of the change in baseline TEPs (** = p < 0.01; *** = p < 0.001), the hash sign shows the significance of the SICI x medication interaction (# = p < 0.05). (C) **MEPs**. MEP SICI before and after the medication is expressed in % of TS MEP (alprazolam: red, placebo: blue). (D-F) **Correlation between the extent of MEP and TEP SICI**. A significant correlation was found in ranked baseline data for SICI in MEPs and SICI in all three modulated TEP components (N17, P60, N100; alprazolam: red, placebo: blue).

**Table 3.** Mean TEP SICI and corresponding statistics. TEP amplitude change was computed as the difference between the CS-TS TEP and the TS TEP, in the table represented in μV ± SEM. Positive numbers mark an increase of the amplitude and vice versa. Displayed statistics correspond to the effect of interest: at baseline we show the effect of stimulus, post medication we show the interaction between SICI and medication.

| peak | time | mean change |  | F statistic | df | p value |
| --- | --- | --- | --- | --- | --- | --- |
|  |  | placebo | alprazolam |  |  |  |
| N17 | baseline | - 3.00 ± 0.84 | - 3.44 ± 0.85 | 39.37 | 1, 58 | < 0.001 *** |
| | post med. | - 3.3 ± 0.95 | - 2.36 ± 0.77 | 3.53 | 1, 54.01 | 0.07 |
| P30 | baseline | - 0.18 ± 0.24 | - 0.78 ± 0.18 | 3.09 | 1, 58 | *n.s.* |
| | post med. | - 0.32 ± 0.3 | - 0.84 ± 0.2 | 0.49 | 1, 54.06 | *n.s.* |
| N45 | baseline | 0.29 ± 0.23 | 0.5 ± 0.22 | 3.52 | 1, 58 | *n.s.* |
| | post med. | 0.41 ± 0.23 | 0.25 ± 0.19 | 0.77 | 1, 54.07 | *n.s.* |
| P60 | baseline | - 1.56 ± 0.49 | - 1.73 ± 0.48 | 12.98 | 1, 58 | < 0.001 *** |
| | post med. | - 1.08 ± 0.29 | - 1.3 ± 0.26 | 0.01 | 1, 54.03 | *n.s.* |
| N100 | baseline | - 1.63 ± 0.81 | - 2.16 ± 0.63 | 10.7 | 1, 58 | < 0.01 ** |
| | post med. | - 1.82 ± 0.54 | - 0.96 ± 0.37 | 5.89 | 1, 54 | < 0.05 * |
| P180 | baseline | 0.17 ± 0.48 | 0.12 ± 0.52 | 0.61 | 1, 58 | *n.s.* |
| | post med. | - 0.14 ± 0.49 | 0.08 ± 0.42 | 0.32 | 1, 54.06 | *n.s.* |

The observed decrease in TEP amplitude was correlated to the decrease of MEPs. Preliminary data visualization revealed a possibly non-linear relationship between TEP and MEP variables, so we calculated Spearman’s correlation coefficients for ranked data that proved significant for all three SICI-sensitive TEP components (Fig 3D-F): N17 (ρ = 0.39, t(38) = 2.64, p < 0.05, adjusted R^2^ = 0.13), P60 (ρ = 0.54, t(38) = 3.96, p < 0.001, adjusted R^2^ = 0.27), N100 (ρ = 0.55, t(38) = 4.08, p < 0.001, adjusted R^2^ = 0.29).

### 3.7. N17 SICI is correlated with changes induced by alprazolam

We postulated that if both the modulation of the N17 component of M1 TEPs by paired-pulse stimulation and the changes observed after alprazolam administration are due to GABA_A_R activation, both should be dependent on interindividual variations in the functional state of GABA_A_R-mediated inhibition in M1. To investigate this further, we evaluated the correlation between the N17 SICI produced by paired-pulse stimulation before medication, and the most prominent changes observed after alprazolam. More precisely, we selected the alprazolam-induced reduction of N17 and the augmentation of the amplitude in the lower β band. A significant linear correlation was observed between N17 SICI and both measures (with the modulation of N17: r(18) = 0.47, t = 2.28, p < 0.05, adjusted R^2^ = 0.18; with the modulation of lower β amplitude: r(18) = -0.61, t = -3.3, p < 0.01, adjusted R^2^ = 0.34; see Fig 4C,D). This indicates that subjects with a larger reduction of N17 following paired pulse stimulation also show a stronger modulation of N17 and β amplitude by alprazolam.

**Fig 4.**
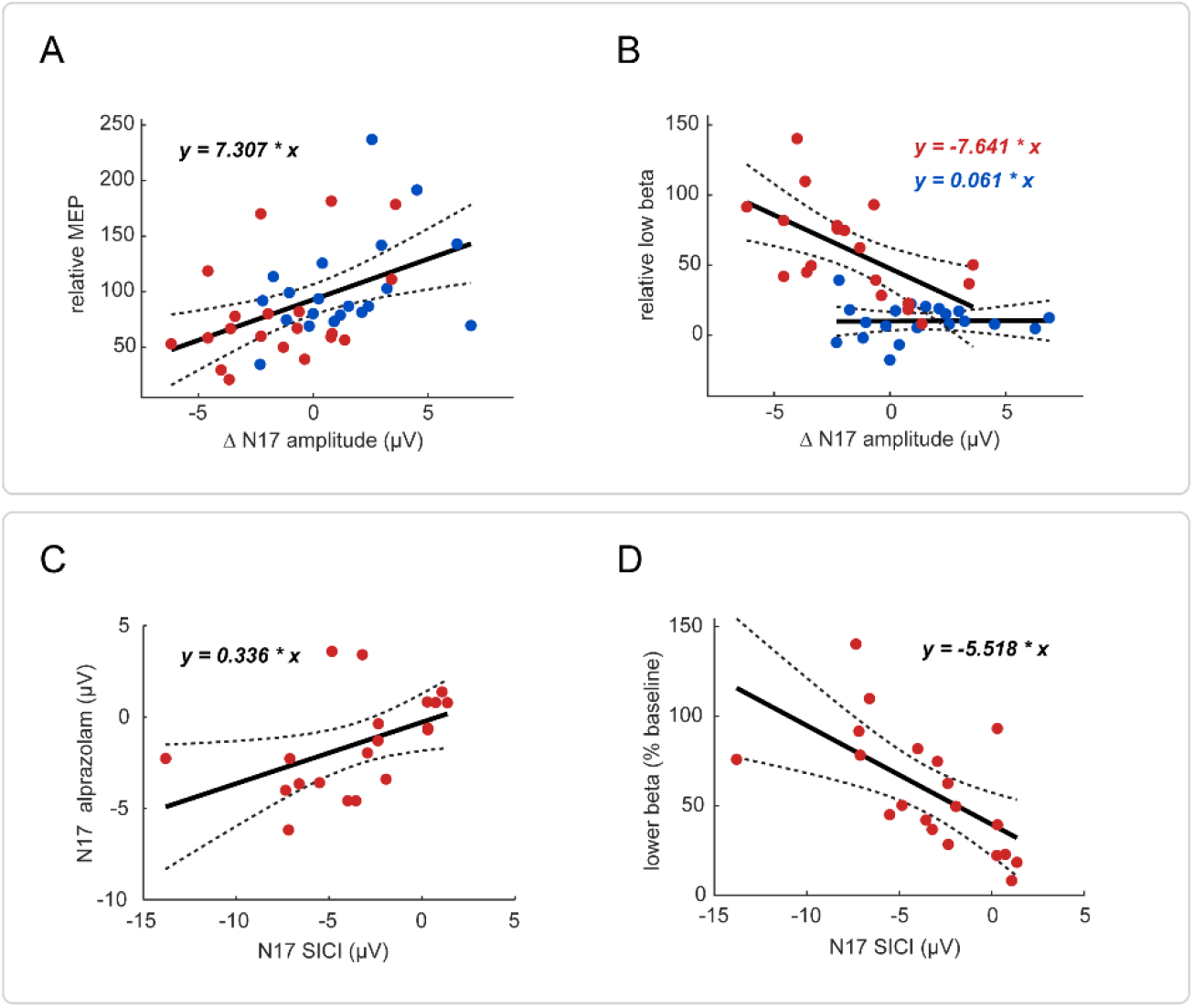
Correlational analysis of TEP component N17. (A) A significant linear correlation was found between the drug-induced modulation of N17 and the drug-induced modulation of MEPs. (B) In the alprazolam session (red), the modulation of N17 was found correlated to the modulation of low β amplitude, whereas no association was found for placebo (blue). The amplitude change of N17 is expressed as a difference from baseline (μV), the change of MEPs and low β as a relative amplitude (% of baseline). (C) The extent of N17 SICI was correlated with the alprazolam-induced modulation of N17 amplitude and (D) of the amplitude of lower β oscillations. N17 measures are expressed as a difference (μV), either from TS TEPs (SICI) or the baseline (alprazolam), low β as a relative amplitude (% of baseline).

### 3.8. SICI tends to be reduced when GABA_A_Rs are potentiated by alprazolam

As a last exploratory analysis, we investigated possible interactions between the effect of paired-pulse stimulation and the effect of alprazolam. There was no significant difference between the MEP reduction induced by paired-pulse stimulation after alprazolam medication (70.93 ± 4.6 %) compared to placebo (73.25 ± 3.97 %) (Fig 3C). Similarly, no significant interaction was found between the type of medication and the effect of paired-pulse stimulation on most TEP components (Fig 3B) with the exception of the N100, whose SICI was reduced by alprazolam and increased by placebo. A tendency in the same direction was also noted in the component N17. Mean observed changes in TEP SICI as well as the outcome of the statistical analysis are summarized in the Table 3.

## 4. Discussion

### 4.1. TEP signature of pharmacological GABA_A_R activation

We investigated the impact of pharmacological activation of GABA_A_Rs on the amplitude of TEPs evoked by stimulation of M1, while also distinguishing M1 TEPs elicited by stimulation at sub- and supra-threshold intensities. Furthermore, three established RS-EEG measures were obtained to further characterize the effect of alprazolam and explore their relation to drug-induced changes in TEPs.

We observed a specific alprazolam-induced modulation of both early and late components of supra-threshold M1 TEPs, whereas only late components were significantly affected in sub-threshold M1 TEPs. The alprazolam-induced modulation of the late N100 and P180 components was robust, consistent across TEP datasets, and clearly reflects changes in brain function resulting from a global increase in GABA_A_R-mediated inhibition. Yet it does not necessarily provide specific information on the state of inhibition in the brain network targeted by TMS. Indeed, there is a growing body of evidence indicating that the N100-P180 complex can be mostly attributed to the brain activity evoked by peripheral sensory stimulation [43, 59-62]. This is compatible with the notion that these late components relate to the vertex potential typically evoked by transient sensory stimuli [63]. The magnitude of this vertex potential can be expected to depend on the excitability of multiple brain networks as well as the state of populations of spinal or peripheral neurons influenced by GABA_A_R-mediated neurotransmission. Indeed, previous studies have shown a benzodiazepine-induced reduction of both auditory- and somatosensory-evoked vertex potentials [64-66]. Therefore, when considering sub-threshold M1 TEPs, the alprazolam-induced modulation of N100 and P180 should not be interpreted as a direct measure of cortical inhibition in the stimulated brain network.

On the other hand, the topography of the late components elicited by supra-threshold M1 TEPs differs from that of sub-threshold M1 TEPs, indicating that additional processes are triggered by supra-threshold stimulation of the motor cortex [43, 67-69], possibly related to the activation of the motor brain network and/or the somatosensory response to the TMS-evoked muscular contraction. The N100 component of supra-threshold M1 TEPs was previously associated with motor inhibition [70-73] that is possibly mediated by local GABA_B_Rs [25]. Therefore, the N100 reduction following administration of alprazolam might reflect both a suppression of unspecific sensory-evoked activity and a decrease in local sensorimotor inhibition following a weaker M1 activation.

The earlier components of sub-threshold M1 TEPs were not significantly affected by alprazolam as compared to placebo. This lack of effect could be related to the fact that these TEP components are relatively small and show substantial inter-subject variability. In contrast, the early components of supra-threshold M1 TEPs were significantly modulated by alprazolam. In line with reports of previous studies exploring the effect of benzodiazepines on TEPs [25, 29], alprazolam was associated with an increase of the N45 component. In past, this component was also found suppressed by a competitive GABA_A_R antagonist selective for α5 receptor subtypes [74], whereas dextromethorphan, an antagonist of the N-methyl-D-aspartate receptor for glutamate (NMDAR), led to its augmentation [75]. Therefore, our results add to the accumulating evidence that N45 depends on the balance between GABA_A_R-mediated inhibition and NMDAR-meditated excitation.

Most remarkably, alprazolam significantly reduced the component N17 of supra-threshold TEPs (labelled N15 or N18 in other studies [76, 77]) when compared to the effect of placebo. This component is of high interest as it likely reflects genuine cortical activity directly evoked by TMS within the stimulated motor network. It is target-specific, seems to be little affected by somatosensory- and auditory-evoked potentials, and its very early latency excludes any contribution of the recurrent somatosensory response to the contraction of the contralateral hand muscle [43, 59, 78]. Furthermore, several previous studies located the sources of N17 to the M1 and/or adjacent nodes of the sensorimotor network [78-82]. Therefore, since our data clearly shows that N17 is dependent on the state of GABA_A_R-mediated neurotransmission, the extent of its modulation may be directly linked to the performance of the GABA_A_ergic inhibitory system within the target brain network. This claim is further supported by the significant correlation between the drug-induced change of the N17 amplitude and the amplitude of MEPs, which confirms the association of this component to the motor output of M1 and hints on its possible relation to the TMS-evoked cortico-spinal activation.

### 4.2. RS-EEG signature of pharmacological GABA_A_R activation

As expected, all three evaluated RS-EEG markers were sensitive to the effects of alprazolam as compared to placebo, the increase of GABA_A_R-mediated inhibition manifesting as a clear augmentation of lower β-band amplitude and reduction of AAC and SE. In addition, the effect of alprazolam on the amplitude of N17 of supra-threshold TEPs was strongly correlated with the alprazolam-induced modulation of the amplitude of lower β oscillations over central regions.

The increase in β represents an established RS-EEG marker of the benzodiazepine effect [45, 47, 83] and while the origin of this emerging activity has not been reliably identified, it is not unlikely that motor networks significantly contribute to its generation. Oscillatory activity in the β band was in the past linked predominantly to the function of sensorimotor cortices [84], where it was proposed to reflect a process maintaining the ongoing postural setting while inhibiting new movements [85]. Increased β activity, either spontaneous or induced by repetitive brain stimulation, was associated with the slowing of voluntary movements [86, 87], which indicates that it might be associated with states of lower motor excitability. Provided that the magnitude of the TEP component N17 reflects local cortical activation of the motor cortex and/or nearby connected areas, our data support the idea that both the alprazolam-induced increase in β oscillations over central regions and the alprazolam-induced decrease of the N17 depend on the functional state of GABA_A_R-mediated inhibition in the motor cortex.

The alprazolam-induced reduction of the N17 was not correlated to the other two RS-EEG markers. The AAC is a measure of the attenuation of α oscillations over occipital regions where α band activity is most pronounced [45]. The SE represents the slope of the 1/f-like background component of the EEG frequency spectrum, is related to the level of arousal and consciousness [54, 88, 89] and is thought to reflect the ratio of global excitatory and inhibitory neurotransmission [90]. The N17 component and both of these RS-EEG markers are thus likely to reflect activity of different brain networks having distinct cytoarchitectural and functional properties. The lack of correlation between the effect of alprazolam on N17 and its effect on the AAC and the SE would indicate that, across individuals, these networks have independent dynamics leading to independent variations in their sensitivity to GABAergic inhibition. Therefore, it might be advantageous to combine TEPs and different RS-EEG measures when evaluating the pharmacologically induced increase in GABA_A_R-mediated inhibition, as they likely provide complementary information.

### 4.3. TEP SICI and its relation to the effect of alprazolam

In the second part of the study, we activated local inhibitory circuits in M1 using a sub-threshold conditioning stimulus producing short-latency intracortical inhibition (SICI). In both baseline recordings, SICI manifested as a significant reduction of MEP amplitudes, and a concomitant amplitude suppression of TEP components N17, P60 and N100. Interestingly, the spatiotemporal distribution of the SICI effect bore a striking resemblance to the PCA component identified by Biabani et al. [78], which allowed to distinguish real M1 TEPs from the brain response evoked by sham stimulation. Assuming that such component can be attributed to the activity directly evoked by TMS in the stimulated cortical area, our findings suggest that the conditioning stimulus indeed led to a suppression of this genuine activity, likely due to a local increase in GABA_A_R-mediated inhibition.

Across individuals, the reduction of MEP amplitude was correlated with the reduction of all three TEP components, corroborating the assumption that changes in the magnitude of these TEP components can be linked to changes in corticospinal motor output. Since we previously identified the N17 as the component most reliably reflecting the changes in the GABA_A_ergic neurotransmission within M1, we further explored its modulation by SICI in relation to its modulation by alprazolam and found, across individuals, a significant linear correlation between the N17 reduction induced by alprazolam and the N17 reduction induced by paired-pulse stimulation. Furthermore, we found a similar relationship between N17 SICI and the alprazolam-induced increase in low β band oscillations. In other words, subjects that showed a stronger effect of pharmacological activation of GABA_A_Rs also showed a stronger effect of SICI, indicating that both phenomena are at least partially mediated by the same mechanism. This further strengthens our assertion that the N17 component can be linked to the functional state of GABA_A_R-mediated inhibition within the motor cortex.

Our observations of TEP SICI are only in partial agreement with previous studies. These mostly reported a suppression of P60 but also found a decrease of P30 [30-32], and a decrease [30] or an increase of N45 [28, 31, 32]. In a very recent publication, Rawji et al. additionally showed a suppression of N15 (here N17) and P180 [28]. Contrary to all above-mentioned studies, the early study of Paus et al. reported no effect of SICI in TEPs [91], whereas two more recent studies only found a reduction of late TEP components N100 and P180/P300 [29, 33]. These discrepancies may be explained by the variability of stimulation parameters as well as the analysis pipeline to extract TEP parameters, which ranged from a point-by-point comparison across all electrodes to peak extraction from electrodes of interest in predefined time intervals varying across studies. Here, for each TEP component, the electrodes of interest were selected based on the mean component topography, to prioritize the most prominent generators while possibly neglecting other sources contributing to the final voltage distribution of TEPs. Furthermore, our analysis technique allowed us to extract the amplitude at subjects’ peak latency and thereby address the issue of inter-subject variability in the TEP temporal profile [92], which is especially important when evaluating early and very transient TEP components.

Finally, we examined whether the local inhibition induced by paired pulse stimulation is affected by the global activation of GABA_A_Rs produced by alprazolam. Contrary to previous studies that reported an increase of SICI in MEPs following the administration of positive GABA_A_R modulators [93-96], we did not observe any significant change of MEP SICI after alprazolam. When considering the influence of alprazolam on TEP SICI, our results are in line with those reported by Premoli et al. who combined paired-pulse stimulation with the administration of diazepam and found a significant reduction of SICI on the N100 component [29]. However, based on the effect of alprazolam we observed in supra-threshold M1 TEPs, we speculate that the observed modulation of N100 SICI might be to a great extent due to the alprazolam-induced suppression of single-pulse N100 amplitude rather than a reduction of the SICI effect itself.

## 5. Conclusion

In the present study, we described the effect of increased GABA_A_R-mediated inhibition on specific components of M1 TEPs using both pharmacological and paired-pulse activation of GABA_A_R inhibitory circuits. In supra-threshold M1 TEPs, we identified an early component N17 that was directly associated with the motor output and was found susceptible to both alprazolam and SICI. The modulation induced by both manipulations was correlated across subjects. In addition, we linked the alprazolam- and SICI-induced reduction of N17 to the increase in lower β band oscillations observed following the administration of alprazolam, indicating that changes in both measures correspond to the increased GABA_A_R inhibitory transmission in the stimulated motor cortex and closely connected areas of the sensorimotor brain network. Taken together, we show that the N17 component of supra-threshold M1 TEPs is sensitive to GABA_A_R-mediated changes in local cortical excitability. Therefore, pharmacological or paired-pulse modulation of this component could be used in clinical settings as a non-invasive biomarker to assess the functional state GABA_A_R neurotransmission in sensorimotor cortices. TEP recordings may be further supplemented by RS-EEG measures (lower β amplitude, AAC, SE) to assess the GABA_A_R inhibition more globally.

## 7. Authors’ contribution

DS: experimental design and methodology, data acquisition, data pre-processing and analysis, visualization, original draft, manuscript editing and finalisation. AS: data acquisition, methodology, manuscript revision. AI: conceptualization, supervision, manuscript revision, funding acquisition. AM: conceptualization, experimental design and methodology, supervision, funding acquisition, manuscript editing and revision.

## 8. Acknowledgments

The research was funded by the ARC grant (“Actions de Recherche Concertées”) provided by Ministère de la Fédération Wallonie-Bruxelles under the project *ARC 17/22-083*, as well as by the Fonds de la Recherche Scientifique (FNRS) of which D. Sulcova is a research fellow.

## 9. Supplementary materials

Supplementary material complementing this manuscript is available online (https://…). The material is divided in four sections: (1) description of TEP pre-processing, with the emphasis on the two-round ICA; (2) introduction to microstate analysis as a source of prior for TEP peak amplitude extraction; (3) description of MEP pre-processing and the identification of epochs with excessive baseline activity; (4) brief description of the effect of alprazolam on TEPs evoked by stimulation of the left angular gyrus.

