## Supplementary materials for "Evaluation of GABA_A_R-mediated inhibition in the human brain using TMS-evoked potentials"

This document provides detailed information about the methodology used for pre-processing and analysis of data presented in a homonymous manuscript submitted to *NeuroImage* in August 2022. The methodology description is divided into three sections: First, we describe the pre-processing pipeline of TMS-evoked potentials (TEPs) with particular emphasis on the Independent Component Analysis (ICA), providing examples of ICA components considered as artifactual and their characteristic features. Second, we introduce microstate analysis of TEPs and demonstrate its utility as a source of prior for peak amplitude extraction. Third, we describe automatized processing of motor-evoked potentials (MEPs) that allows to identify and exclude trials with excessive baseline activity while minimizing subjective input.

In the last section of this document, we present TMS-evoked potentials obtained by stimulation of the left angular gyrus (AG) that were recorded in addition to TEPs from the primary motor cortex (M1), motor-evoked potentials and resting-state EEG. Similarly to these measures, we evaluated the effect of alprazolam on AG TEPs and the results of the analysis are presented here. Since this part of the study does not significantly contribute to the overall conclusions we make, for the sake of clarity it was not included in the main publication.

#### Pre-processing of TMS-evoked potentials

Three types of TEPs (conditions) were investigated in the present study: TEPs obtained by supra-threshold stimulation of the left primary motor cortex (M1), sub-threshold M1 stimulation, and stimulation of the left angular gyrus (AG). 2 TEP datasets of each condition were obtained in each of the two experimental sessions, one at the baseline and the other following the medication (alprazolam or placebo). All TEPs were recorded using the NeurOne system (Bittium) with 32 open-circle electrodes (Multitrodes, EasyCap), sampling rate of 20 kHz and mastoid electrodes serving as reference.

The raw signal was pre-processed in several automatized steps:

1. Signal was re-referenced to Cz, mastoids were removed, and the signal was re-referenced again to the average calculated from 30 remaining electrodes
2. DC shift was removed from the whole recording
3. Signal was segmented into epochs relative to TMS stimulus [-1 2]s
4. TMS artifact was cut out between [-0.005 0.01]s and replaced using cubic interpolation
5. Signal was down-sampled by 10, so the size of a single epoch corresponded to 6000 bins

6. Linear trend was filtered out from individual TEP epochs
7. Single trials were split into appropriate categories, missed trials identified during the recording session were removed
8. Data were visually checked for bad channels that were removed and interpolated using the average signal of the 6 surrounding electrodes (this happened for 1 channel in 3 datasets out of 40)

*First round of ICA.* In the next step, we calculated a square matrix of 30 independent components (ICs), unless a channel was previously interpolated – in that case we lowered the number of components by one. The ICs were visually inspected, and 1-2 ICs (rarely up to 4) were identified as containing the artifact caused by muscular contraction due to superficial TMS stimulation. Such artifact can be easily spotted in the single-trial time-course as a high-voltage deflection with a decay tail and a sharp peak at 10 ms, which corresponds to the limit of the interpolated interval. It shows typical lateralized topography that varies across stimulation sites as well as across subjects according to their anatomy (Figure 1A).

Identified components were removed, and new signal was created based on the remaining ICs. It was then filtered with a Butterworth bandpass filter ($4^{th}$ order) to remove extreme frequencies, keeping values between 0.1 and 80 Hz. The line noise at 50 Hz was removed with a FFT notch filter cutting out frequencies between 48-52 Hz with a 2 Hz wide Hanning function. Next, the epochs were cropped from [-0.75 0.75]s relative to the stimulus in order to remove possible distortion caused by frequency filters. Finally, each dataset was visually inspected trial-by-trial and epochs that included isolated bursts of high-amplitude activity, extreme drifts etc. were discarded.

*Second round of ICA.* In the second round, we calculated a rectangular ICA matrix for the total number of channels (30) minus the number of ICs removed in the previous round of ICA. We also extracted specific features of each IC, such as its average time-course and the frequency spectrum up to 45 Hz. As a preliminary step, we used Multiple Artifact Rejection Algorithm (MARA) [1], a linear classifier originally trained on adult EEG data, to identify components possibly containing an artifact. Selected ICs were then individually inspected by an experienced researcher. Based on their features, these ICs were classified either as *mixed* (containing signal of likely cerebral origin) and retained, or assigned to one of the following categories and removed: (1) eye movements such as blinking and saccades (Figure 1B); (2) tonic muscular activity (Figure 1C); (3) decay artifact (Figure 1D); (4) electrode noise (Figure 1E); (5) ringing artifact introduced by frequency filters (Figure 1F).

*Figure 1*

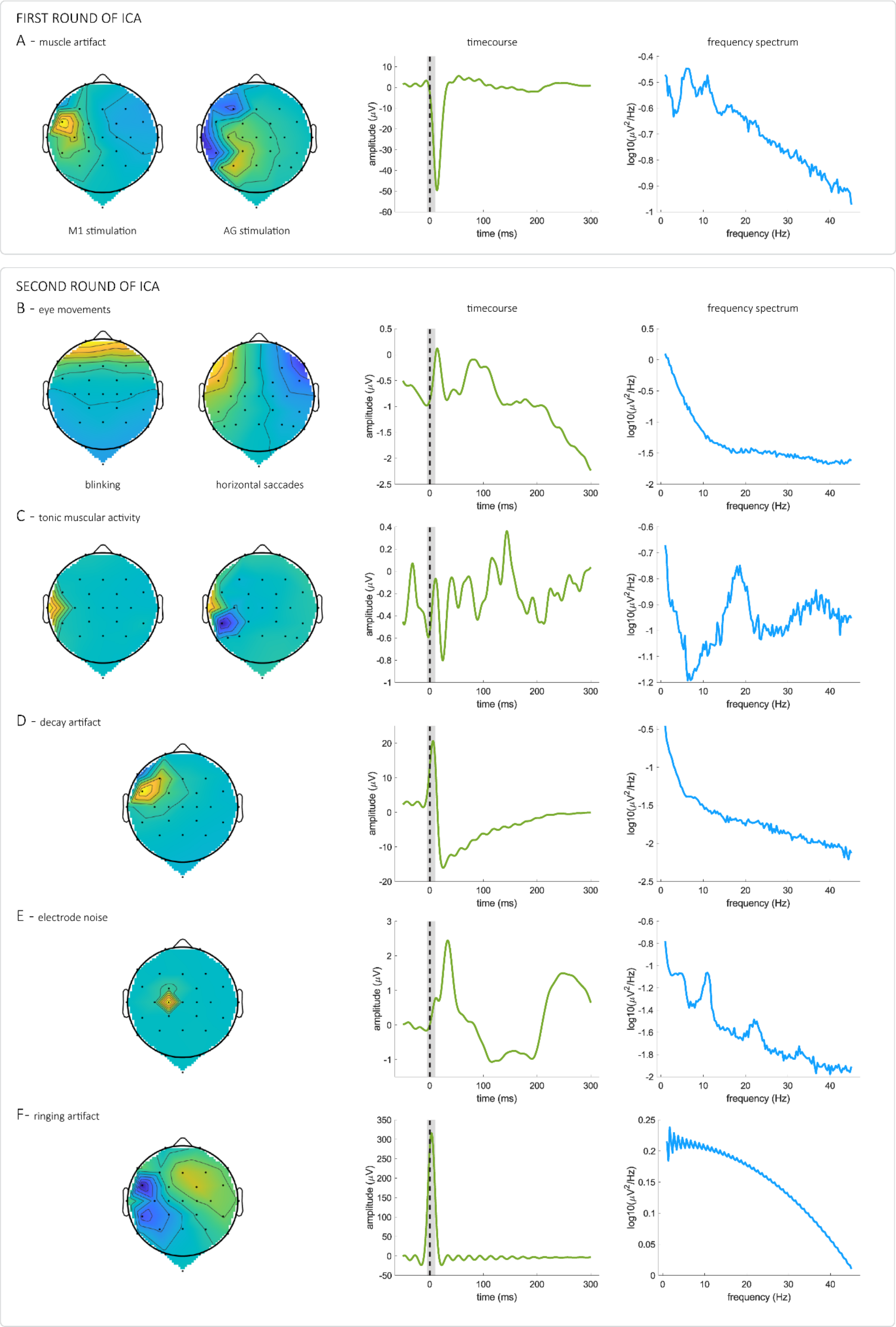

FIRST ROUND OF ICA
A - muscle artifact
M1 stimulation
AG stimulation
timecourse
frequency spectrum
amplitude (μV)
time (ms)
log10(μV²/Hz)
frequency (Hz)
SECOND ROUND OF ICA
B - eye movements
blinking
horizontal saccades
C - tonic muscular activity
D - decay artifact
E - electrode noise
F- ringing artifact

Final TEP epochs, composed of the signal of remaining ICs, were then baseline corrected by subtracting from each timepoint the average amplitude of [-0.2 -0.005]s relative to the stimulus and averaged, thus resulting in a single TEP waveform per subject per condition.

**Microstate analysis as a source of prior for TEP peak amplitude extraction**

All TEPs were evaluated between [0.01 0.3]s post stimulus. In the first step, baseline TEPs of each category were characterized using microstate analysis, a method that considers the temporal evolution of the whole scalp topography and identifies discrete time intervals when a single topographic pattern dominates [2]. It relies on the assumption that two different distributions of voltage in the same dataset represent the activation of two distinct sets of cortical sources [3]. In practice, all timepoints in a group average dataset are subjected to a clustering process, which defines a number of microstate classes and labels temporally adjacent timepoints with the most fitting class, thereby forming non-overlapping microstate intervals [4]. These segments closely match observed TEP components and in evoked potentials they were suggested to represent a sequence of discrete processing steps triggered by the stimulation [2]. Importantly, microstate analysis evaluates TEPs normalized by their global field power (GFP, see Equation 1), i.e. the mean amplitude across all electrodes, thus limiting the effect of the inter-subject variability in the response strength. Moreover, by considering the whole voltage landscape, the method deals with the bias introduced by the choice of a reference electrode.

*Equation 1*

$$GFP(t) = \sqrt{\frac{\sum_{j=1}^{n}(u_j(t) - \bar{u}(t))^2}{n}}$$

To perform the microstate analysis, we used the open-access Matlab toolbox *Ragu* and followed the recommended pipeline [5]. For each TEP condition at the baseline, we obtained a sequence of microstate class maps that were associated with individual TEP components (a detailed topographic analysis of baseline TEPs and their comparison was previously published in a separate article [6]). We then used spatiotemporal features of these microstates to inform the subsequent semi-automatic extraction of TEP peak amplitude (Figure 2). The duration of each microstate provided a base for the calculation of the default width of the extraction window that was kept identical across subjects and tested datasets. The topographic pattern of the microstate was used to identify three electrodes of interest (EOIs) that corresponded to the spatial maximum of the associated TEP component. The signal of these EOIs was averaged and all compared datasets from one subject/condition were plotted together within the exploration window. The microstate latency was used as the starting window latency that was adjusted manually to fit TEP peaks of the individual subject, thus accounting for the substantial inter-subject variability of TEP temporal profile. Finally, local amplitude maxima of all compared datasets (or minima, in case of negative components) were automatically measured in the exploration window. All above mentioned steps were performed using in-house written Matlab scripts.

Using this approach, we extracted a single value for each subject, tested dataset and TEP component, thereby substantially reducing the number of comparisons necessary for the statistical analysis. We chose to evaluate the significance of amplitude changes for each TEP component separately using a linear mixed model that allowed individual variability in the intercept but obtained data could have been assessed by any other classic model. Figure 3 provides an overview of microstate topographies and identified EOIs for all components of each evaluated TEP condition.

*Figure 2*

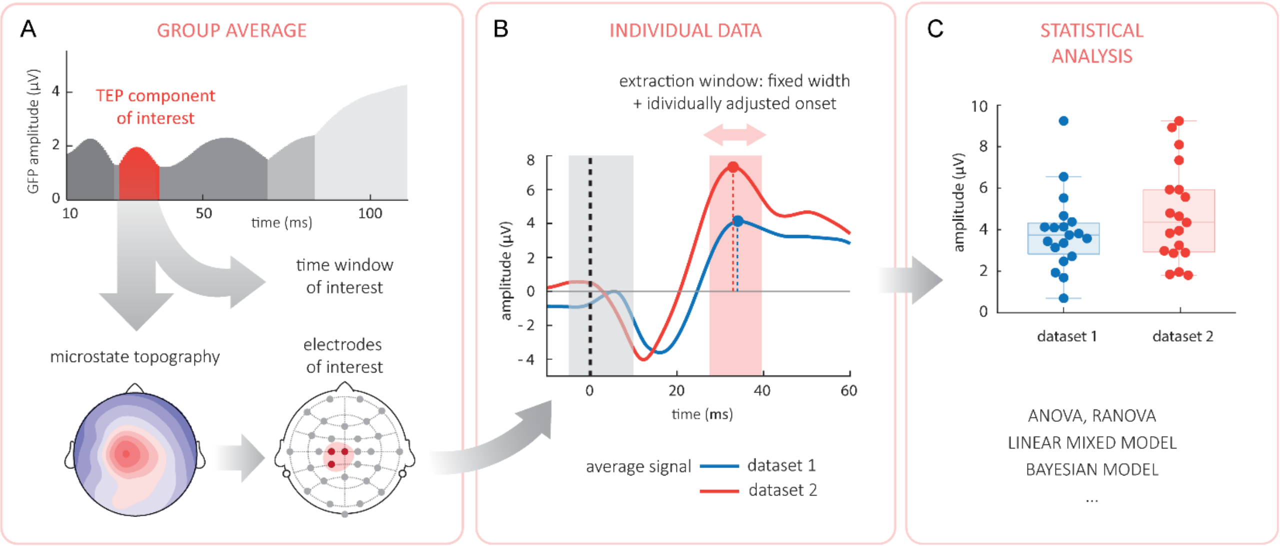


**TEP peak amplitude extraction.** A - the result of clustering that attributed one microstate class to each TEP component (visualized as distinct peaks in the GFP of the average signal). Mean microstate topography was used to identify EOIs, latency and the width of the exploration window attributed to the evaluated component. B - the interactive plot displaying at the same time the averaged EOI signal of all compared datasets of a single subject. The onset of the exploration window is manually shifted to include all local peaks. Thereby extracted values can be then included into any type of classic statistical analysis, as visualized in the panel C.

*Figure 3*

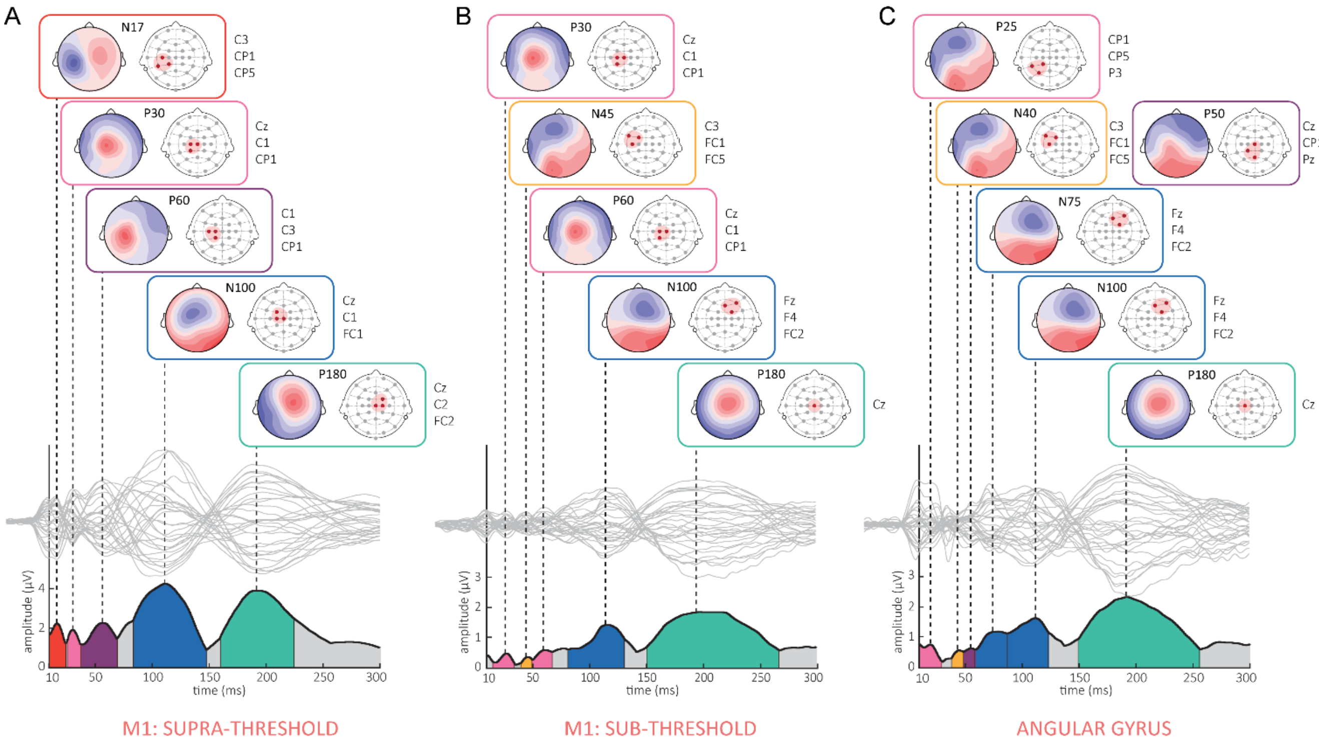


**Microstate topographies and EOIs of TEP components.** Each vertical panel corresponds to one TEP condition: A - supra-threshold M1 TEPs; B - sub-threshold M1 TEPs; C - AG TEPs. In the bottom part, the GFP of the average baseline TEP is shown and the duration of microstates assigned to TEP components is shaded in corresponding colors. In the middle, average baseline signal from individual electrodes is plotted in light gray and dashed vertical lines mark peak latencies of TEP components. In the upper part, microstate class maps are linked to each TEP components and EOIs are visualized at the EEG layout, labels displayed on the side. N45 in supra-threshold M1 TEPs was not identified therefore the spatiotemporal features of N45 in sub-threshold M1 TEPs were used for the amplitude extraction.

#### Pre-processing of motor-evoked potentials

MEPs were recorded using an EMG system connected to the neuronavigation (TMSi MOBI, sampling rate 1024Hz) in parallel to TEPs when delivering the supra-threshold stimulus over M1. Raw EMG signals were automatically pre-processed in the following steps:

1. Signal was corrected for DC shift
2. Frequencies below 4 Hz were filtered out with a Butterworth high-pass filter ($4^{th}$ order)
3. Signal was segmented into epochs relative to TMS stimulus [-0.2 0.4]s
4. Linear detrend was applied to individual epochs
5. Single trials were split into appropriate categories, missed trials identified during the recording session were removed

In the next step, we identified and discarded trials with excessive baseline activity, as it might indicate a state of increased excitability, possibly of sub-cortical origin. To this end, we used an in-house written Matlab script that automatized this procedure, thereby minimizing the subjective input. The cleaning comprised two distinct steps. First, a root mean square (RMS) value of the baseline was calculated for each trial and epochs whose RMS exceeded the value of mean RMS ± 3 SD were discarded. The process was then repeated in remaining epochs until no outliers were identified. Second, an arbitrary threshold was selected based on normally observed EMG noise (we set it at ± 15µV) and epochs showing larger amplitude at the baseline were also excluded. An output figure obtained after the cleaning of a single dataset is shown in Figure 4.

Finally, we baseline-corrected the signal by subtracting the mean amplitude of the interval between [-0.2 0]s relative to the stimulus and extracted the peak-to-peak MEP amplitude at single trial level. Mean values were calculated for each subject/tested dataset and included in the statistical analysis.

*Figure 4*

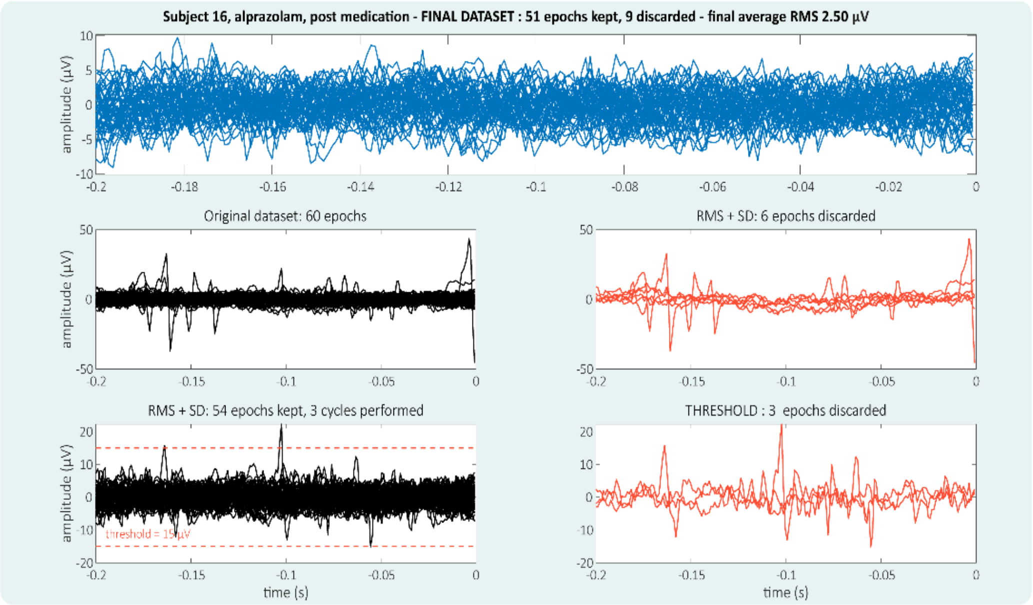


**Example of the output figure of the MEP cleaning process.**

### TMS-evoked potentials from the angular gyrus

*AG TEPs.* The stimulation target in the left AG was determined in the individual MRI using the position of the superior temporal sulcus (STS) and the intraparietal sulcus (Figure 5A). Using neuronavigation, the coil was positioned with the handle pointing downwards and 10° towards the midline, thus placing the coil axis perpendicularly to the STS and across the AG (Figure 5B). 60 TMS stimuli of 100% rMT were delivered over the target in a single block with a variable inter-trial interval (4– 6 s) while simultaneously recording EEG. At the baseline (before any medication), AG TEPs had a distinct spatiotemporal profile composed of early components P25, N40, P50 and N75 followed by late components N100 and P180 (Figure 5C). While the early components were clearly different from those observed in M1 TEPs, the N100-P180 complex showed features generalized across target sites (for a detailed analysis of baseline AG TEPs see [6]).

*The effect of alprazolam*. The peak amplitude of individual TEP components served as the main outcome variable when evaluating the effect of medication on AG TEPs. Mean observed changes and the outcome of the statistical analysis are summarized in Table 1. Like in sub-threshold M1 TEPs, only late components N100 and P180 of AG TEPs were clearly suppressed by alprazolam as compared to placebo, while earlier components showed no significant change in amplitude (Figure 5D-E).

*Table 1*

| stimulus | peak | mean change in amplitude | | F statistic | df | p value | |
|---|---|---|---|---|---|---|---|
| | | placebo | alprazolam | | | | |
| | **P25** | - 0.30 ± 0.36 | 0.31 ± 0.36 | 1.40 | 1, 18.54 | *n.s.* | |
| | **N40** | - 0.31 ± 0.19 | 0.02 ± 0.27 | 1.02 | 1, 18.54 | *n.s.* | |
| **AG** | **P50** | - 0.13 ± 0.23 | - 0.69 ± 0.26 | 2.64 | 1, 18.54 | *n.s.* | |
| **100 %rMT** | **N75** | - 0.35 ± 0.42 | - 0.14 ± 0.35 | 0.17 | 1, 18.47 | *n.s.* | |
| | **N100** | - 0.70 ± 0.36 | - 1.74 ± 0.46 | 5.07 | 1, 18.31 | < 0.05 | * |
| | **P180** | - 0.20 ± 0.47 | - 3.18 ± 0.73 | 12.76 | 1, 18.54 | < 0.01 | ** |

**Mean change in TEP amplitude and corresponding statistics**. Amplitude change was computed as the difference between measurement post-medication and the baseline, in the table represented in μV ± SEM. Positive numbers mark an increase of the amplitude and vice versa, in case of negative components original voltage measures were multiplied by -1.

*Figure 5*

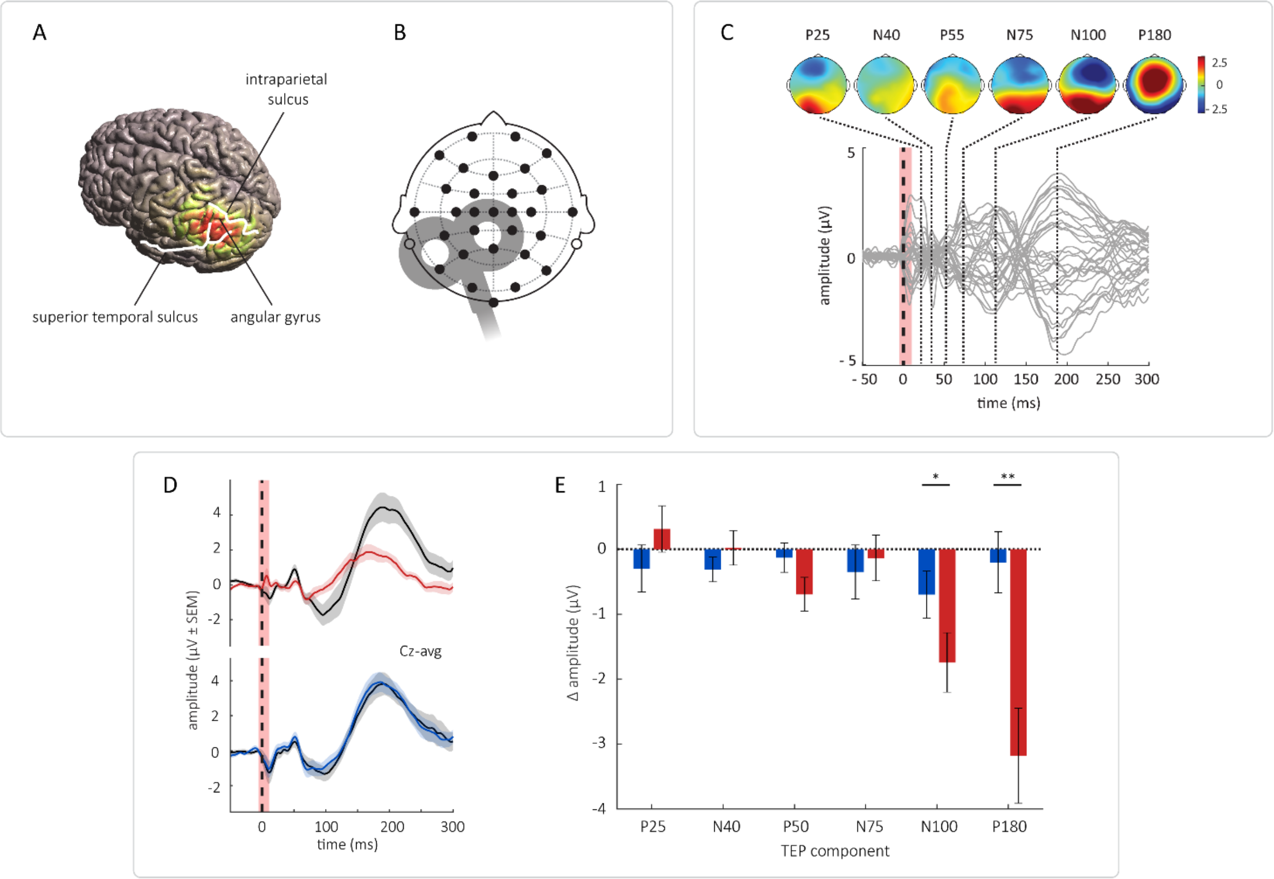


**AG TEPs.** A – a brain model showing the spread of evoked magnetic field across the cortex when stimulating the left AG. Anatomical landmarks are highlighted in white. B – an approximate placement of the TMS coil over the head showing the EEG layout. C – AG TEPs in a baseline recording of one experimental session. Grey traces represent the signal of all electrodes recorded at the baseline in the placebo sessions (there were no substantial differences between baseline TEPs of both sessions), TMS stimulus is marked by thick black dashed line, interval interpolated following the removal of TMS artifact is shaded in pink. Dotted black lines mark peak latencies of TEP components, associated topographies are shown in the upper part of each figure. D - group average TEPs before and after the medication from the most representative electrode (Cz). Baseline: black, post alprazolam: red, post placebo: blue. Shaded areas represent the SEM. E - modulation of peak TEP amplitude. Bar height corresponds to the mean change ± SEM. Positive numbers mark an increase of the amplitude and vice versa, in case of negative components the original voltage measures were flipped to better convey the change induced by the medication. Asterisks denote the significance of the difference between the two sessions (* = $p < 0.05$; ** = $p < 0.01$).
